# The bacterial Bmt methionine synthase is involved in lag phase shortening

**DOI:** 10.1101/2024.06.19.599700

**Authors:** Delia A. Narváez-Barragán, Martin Sperfeld, Einat Segev

**Author notes:** These authors contributed equally to this work (Delia A. Narváez-Barragán, Martin Sperfeld). Institute of Microbiology, ETH, Zurich, 8093, Switzerland.

## Abstract

Bacteria can shorten their lag phase by utilizing methyl groups from compounds such as dimethylsulfoniopropionate (DMSP). These methyl groups are then incorporated into cellular building blocks via the methionine cycle. However, the specific contribution of bacterial methionine synthesis, which is critical for assimilating and incorporating methyl groups, remains unclear.

In this study, we employed transcriptomics, genetic manipulation and biochemical assays to explore the involvement of methionine synthesis in lag phase shortening using the model marine bacterium *Phaeobacter inhibens*. We mapped the expression profiles of the MetH-like methionine synthase components—an enzyme complex that is encoded by three genes—in response to DMSP during the lag phase. Our findings revealed transcriptional decoupling of the three genes. The deletion of the homocysteine-binding component of the MetH-like complex, namely *bmt*, disrupted lag phase shortening in response to DMSP. Through heterologous expression of the *bmt* gene product, we show that the individual Bmt enzyme produces methionine by directly demethylating DMSP and betaine *in vitro*. These findings reveal a metabolic route that was not previously described in marine bacteria. Since Bmt does not require tetrahydrofolate or cobalamin as co-factors for methionine synthesis, its potential to act alone as a demethylase and a methionine synthase represents a cost-effective metabolic shortcut for methyl group assimilation, which could be specifically beneficial under limiting conditions. Indeed, we show that under stress conditions, Bmt allows cells to shorten their lag phase in response to DMSP.

This study enhances our understanding of the enzymatic mechanisms underlying bacterial lag phase shortening, revealing microbial adaptation strategies in response to environmental conditions.

## Introduction

The lag phase describes the transition from dormancy to cell division^1^. Bacteria are under constant evolutionary pressure to optimize their lag phase^2^, which can be achieved by using substrates from the environment to overcome biosynthetic bottlenecks^3,4^. Short lag phases allow cells to rapidly respond to changing environmental conditions, which can be advantageous in dynamic ecosystems. The methionine cycle is a central metabolic pathway involved in regulating the length of the lag phase^3^. Bacteria can accelerate their lag phase by utilizing methyl groups from external methylated donor compounds, such as dimethylsulfoniopropionate (DMSP) and glycine betaine, and incorporate them into building blocks via the methionine cycle^3^. This cyclic pathway is responsible for the synthesis of the amino acid methionine^5^, which is also essential for protein translation initiation^6^ and various biosynthetic pathways^7^. Methionine synthesis is widely distributed across all domains of life, underscoring its importance in cellular metabolism, while the involved enzymes and regulatory elements exhibit evolutionary plasticity^8^.

Methionine synthases play a crucial role in the methionine cycle, catalyzing the methylation of homocysteine to methionine^5^. Methionine synthases can be grouped based on their co-factor requirements (Fig. 1A). The cobalamin-dependent methionine synthase—MetH^9^—is a large multi-domain complex that catalyzes the sequential transfer of a methyl group from methyltetrahydrofolate (CH_3_-THF) via cobalamin (B_12_) to homocysteine (Fig. 1A, reaction 1). In contrast, the cobalamin-independent methionine synthase—MetE—can directly transfer a methyl group from CH_3_-THF to homocysteine (Fig. 1A, reaction 2), but it functions with reduced rates^10^. Minimal versions of MetE were also described, named core-MetE, which lack the tetrahydrofolate-binding domain^11^ and that often utilize methylcobalamin (CH_3_-B_12_) as a methyl group donor (Fig. 1A, reaction 3)^12^. Finally, *N*- and *S*-methylated substrates such as betaine^13^ or *S*-methylmethionine (SMM)^14^ can serve as direct methyl group donors for homocysteine methylation, without involving intermediate methyl group-carrying co-factors (Fig. 1A, reaction 4). The latter methionine synthesis route is catalyzed by minimal versions of MetH, lacking the cobalamin and tetrahydrofolate-binding domains. These simplified versions are designated Bmt (betaine methyltransferase^13^) in the current study. The diversity of methionine synthases, which occurs across all domains of life, highlights the evolutionary adaptations and the importance of this enzyme in the methionine cycle^8^. The specific role of bacterial methionine synthases in the process of lag phase reduction remains unclear.

Here, we explore the involvement of methionine synthases in lag phase reduction of the model marine bacterium *Phaeobacter inhibens*. This bacterium was described to possess a split methionine synthase, in which the domains of the MetH-like methionine synthase are separated into single proteins^15^. It was suggested that all enzymes together build the canonical MetH complex, which produces methionine from CH_3_-THF (which is produced intercellularly from e.g. serine; Fig. 1A, reaction 1). However, considering that *P. inhibens* is a marine bacterium that is often associated with phytoplankton partners^16–18^, the presence of a metabolic shortcut is possible as in the ocean, phytoplankton-derived *N*- and *S*-methylated compounds such as betaine or DMSP are abundant^19,20^. In the presence of these compounds, Bmt has the potential to act as a stand-alone enzyme that transfers methyl groups directly from external betaine or DMSP to homocysteine, producing methionine without involving the co-factors tetrahydrofolate or cobalamin (Fig. 1A, reaction 4). Furthermore, the *P. inhibens* genome encodes a MetE methionine synthase, which may utilize phytoplankton-derived methylated compounds for methionine synthesis (Fig. 1A, reaction 2) while involving additional THF-dependent enzymes such as the DMSP demethylase (DmdA)^21^.

In this study, we employed transcriptomics, genetic manipulation, and biochemical assays to assess the transcriptional response and the involvement of the individual methionine synthase components of *P. inhibens* during DMSP-induced lag phase shortening^3^. Our findings reveal that the Bmt methionine synthase plays a central role in lag phase shortening and can function as a stand-alone methionine synthase that demethylates DMSP and betaine for direct homocysteine methylation. Bmt might therefore represent an alternative route for the assimilation of methyl groups from DMSP and betaine, particularly under unfavorable conditions. This study contributes to our understanding of the enzymatic mechanisms underlying the bacterial ability to shorten the lag phase and underscores the centrality and flexibility of bacterial methionine synthesis.

## Results

### Regulatory dynamics of methionine synthase genes in response to DMSP during the lag phase

We first examined the transcriptional expression patterns of genes putatively involved in methionine synthesis during the lag phase once stationary bacteria are transferred to a fresh medium. Cultures were sampled during the first 15 and 40 min of the lag phase, after bacteria were transferred to fresh medium containing 1 mM glucose as the sole carbon source. Cultures were either supplemented with DMSP (50 µM) or untreated as control. As outlined in the introduction, it was previously shown that DMSP-dependent methionine synthesis in *P. inhibens* involves a MetH-like split methionine synthase complex and proceeds via the sequential transfer of a methyl group from DMSP to tetrahydrofolate (THF)^22^ (*dmdA*), then to cobalamin (*PGA1_c16040*)—involving a cobalamin-binding protein (*cbp*)—and finally to homocysteine (*bmt*), resulting in methionine (Fig. 1B)^15,23^. Our RNA-sequencing data showed that the transcription levels of the three split methionine synthase genes were variable, showing no apparent co-regulation (Fig. 1B, S1 and Table S1). Observed expression patterns ranged from DMSP-induced upregulation (*cbp*) to no regulation (*PGA1_c16040*) and downregulation (*bmt*). The key gene for methionine synthesis (*bmt*) was significantly downregulated in response to DMSP at both examined timepoints. At first look, *bmt* downregulation appears contradictory to the involvement of Bmt in DMSP methyl group assimilation. However, it is in accordance with reports on negative feedback regulation of methionine cycle components in response to their own products^7,24,25^. The downregulation of *bmt* could thus prevent the accumulation of inhibitory methionine, which we previously showed extends the lag phase of *P. inhibens*^3^.

**Figure 1.**
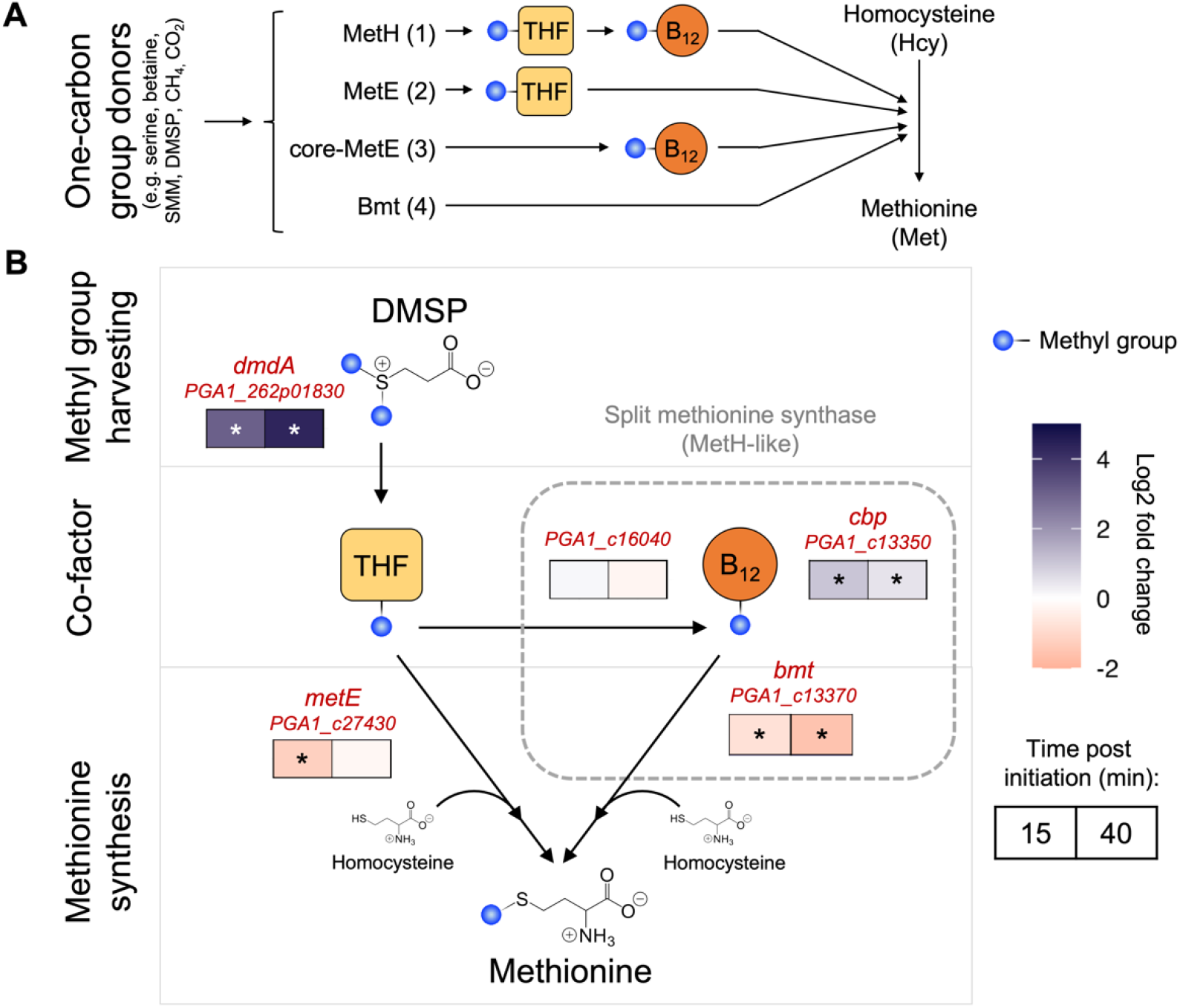
Transcriptional response of demethylation and methionine synthase genes in *P. inhibens* bacteria elicited by DMSP during the lag phase. **(A)** Schematic overview of methionine synthesis reactions, which can be grouped based on the involvement of the co-factors tetrahydrofolate (THF) or cobalamin (B_12_). Numbers in brackets refer to enzymatic reactions described in the main text. **(B)** The transcriptional response of genes putatively involved in *P. inhibens* methionine synthesis was analyzed in freshly initiated bacterial cultures during the lag phase. Glucose-grown stationary phase bacteria were transferred to fresh medium containing 1 mM glucose and supplemented with 50 µM DMSP and were compared to control cultures without DMSP. Changes in gene expression (log2 fold change) are indicated as colored boxes that correspond to the time post initiation—15 min (left box) and 40 min (right box). Colors show upregulation (purple) and downregulation (orange) in response to DMSP. The gene involved in DMSP demethylation (*dmdA*) was highly upregulated in response to DMSP. The three genes encoding the split methionine synthase exhibited a combination of upregulation (*cbp*), no regulation (*PGA1_c16040*) and downregulation (*bmt*). The cobalamin-independent methionine synthase *metE* was downregulated 15 min post initiation. Asterisks indicate significant differential gene expression (adjusted *p*-value < 0.05; log2 fold change < -0.585 and > +0.585). The underlying dataset was previously described in Sperfeld *et al*.^3^. *PGA1_c16040*: methyltetrahydrofolate—cobalamin methyltransferase; *cbp*: cobalamin-binding protein; *bmt*: betaine methyltransferase.

### The methionine synthase Bmt is involved in lag phase shortening in response to DMSP

To further investigate the mechanism by which DMSP methyl groups are utilized for methionine synthesis, we wished to evaluate whether Bmt plays a role in DMSP-driven lag phase shortening in *P. inhibens* bacteria. Therefore, we deleted the *bmt* gene and analyzed bacterial growth and lag phase shortening capabilities. The *Δbmt* mutant is an auxotroph for methionine and therefore growth experiments were initiated with bacteria that were pre-cultivated with 1 mM glucose and 200 µM methionine until reaching stationary phase. Indeed, methionine was required to restore growth of the *Δbmt* mutant (Fig. 2A). Under both conditions, with and without methionine, DMSP did not induce shorter lag times in the *Δbmt* mutant (Fig. 2A). This suggests that Bmt is involved in the lag phase shortening response towards DMSP.

Of note, residual growth was observed in the *Δbmt* mutant in the absence of methionine (Fig. 2A). This growth might result from the presence of a second methionine synthase, namely MetE^26^ (Fig. 1B). The MetE enzyme, which uses CH_3_-THF as a methyl group donor for cobalamin-independent methionine synthesis, could partially compensate for the missing Bmt^27^. Therefore, to evaluate whether MetE is involved in lag phase shortening, we generated a *ΔmetE* mutant. The mutant exhibited normal lag phase shortening upon supplementation with 2 µM DMSP (Fig. 2B). Thus, the methionine synthase Bmt, but not MetE, is involved in lag phase shortening in *P. inhibens* bacteria.

**Figure 2.**
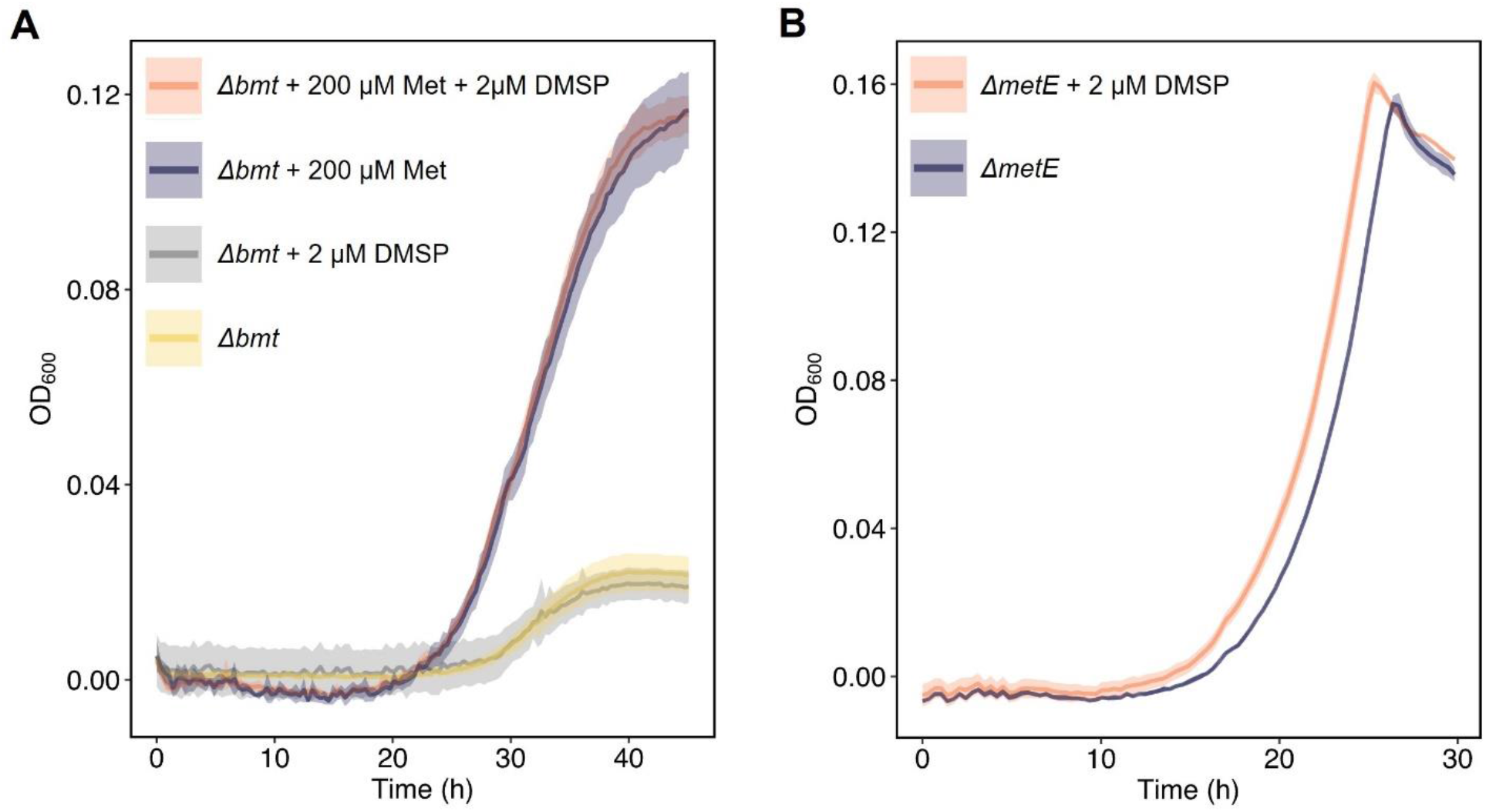
The *P. inhibens* methionine synthase gene *bmt* is required for lag phase shortening. **(A)** Growth of the Δ*bmt* mutant on 1 mM glucose was impaired in the absence of methionine (yellow) but was restored upon adding 200 μM methionine (Met) to the medium (purple). The supplementation of bacteria with 2 µM DMSP did not affect the lag time of the Δ*bmt* mutant, neither in the presence of methionine (orange), nor in its absence (gray). **(B)** The *P. inhibens* methionine synthase *metE*, which encodes for an enzyme that utilizes methyltetrahydrofolate as methyl group donor, is not involved in lag phase shortening. The Δ*metE* mutant still exhibited lag phase shortening upon supplementation with 2 μM DMSP (orange), compared to control cultures (purple). Lines represent the average growth curve based on three biological replicates and shaded areas indicate the standard deviation (SD).

### Bmt acts in vitro as a co-factor independent methionine synthase

Bmt is involved in shortening the lag phase (Fig. 2A), however, it is still unknown if the enzyme functions together with the other MetH-like split methionine synthase components (Fig. 1B), or if it can act as an individual enzyme. Indeed, it was previously shown that homologues of the enzyme Bmt can transfer methyl groups from a variety of methylated compounds directly to homocysteine, without involving THF or cobalamin as methyl group carriers. For example, the Bmt homolog in human, named BHMT, uses a methyl group from *N*-methylated betaine or *S*-methylated sulfobetaine (DMSA) for methionine synthesis^28,29^. A similar substrate spectrum was reported for the Bmt of the bacterium *Sinorhizobium meliloti*^13^. The Bmt homolog in *E. coli* (MmuM^14,30^) and the BHMT-2 enzyme in human^31^ can both transfer a methyl group directly from *S*-methylmethionine (SMM) to homocysteine. Furthermore, it was reported that mammalian Bmt homologues utilize DMSP as a methyl group donor^32,33^. Additionally, transcripts of *bhmt* (*bmt*) from the abundant marine bacteria SAR11 were detected in environmental samples, despite the absence of genes that encode for both the MetE and the MetH enzymes in these bacteria, suggesting the direct utilization of methyl groups from *N*-or *S*-methylated compounds for methionine synthesis *in situ*^34^. These previous reports supported the possibility for direct DMSP demethylation coupled to methionine synthesis by Bmt in *P. inhibens*. Such a direct route for demethylation and methionine synthesis that bypasses the need for the DMSP demethylase DmdA and the co-factors tetatrahydrofolate or cobalamin (Fig. 1B), would represent a novel route for DMSP demethylation in marine bacteria.

To determine whether the Bmt of *P. inhibens* can transfer methyl groups from DMSP directly to homocysteine, we conducted an *in vitro* assay using the purified enzyme. Bmt was expressed in a heterologous system and purified by affinity chromatography (fig. S2). The Bmt enzyme was then incubated with either DMSP or betaine as methyl group donors, and homocysteine as a methyl group acceptor. Purified *E. coli* MmuM was used as a positive control with SMM as a methyl group donor^14^. A commercial fluorescence-based kit was used to measure methionine formation. Our results show that Bmt indeed produced methionine *in vitro* by transferring a methyl group directly from DMSP to homocysteine (Fig. 3). The Bmt enzyme of *P. inhibens* also utilized betaine as a methyl group donor (Fig. 3), a finding of potential significance considering the limited understanding of betaine demethylases in *P. inhibens*. Thus, *in vitro*, Bmt functions as both a DMSP—homocysteine *S*-methyltransferase and a betaine— homocysteine *N*-methyltransferase. Our results do not exclude the possibility that Bmt is part of a MetH-like split methionine synthase *in vivo*^15^, however, the results highlight that the individual Bmt enzyme has the potential to demethylate DMSP and betaine in marine bacteria.

It is important to note however that *in vivo*, Bmt does not appear to replace the key demethylase DmdA in the demethylation of DMSP. This conclusion can be drawn from our previous results with a *ΔdmdA* mutant^3^. In this mutant, the DMSP-dependent lag phase shortening was abolished despite the presence of a *bmt* gene. Taken together, our data demonstrate that the *P. inhibens* Bmt can act *in vitro* as a stand-alone methionine synthase, using different methylated compounds as methyl group donors. These findings underscore the potential flexibility of the enzymatic machinery responsible for harvesting and assimilating methyl groups from diverse methylated donors.

**Figure 3.**
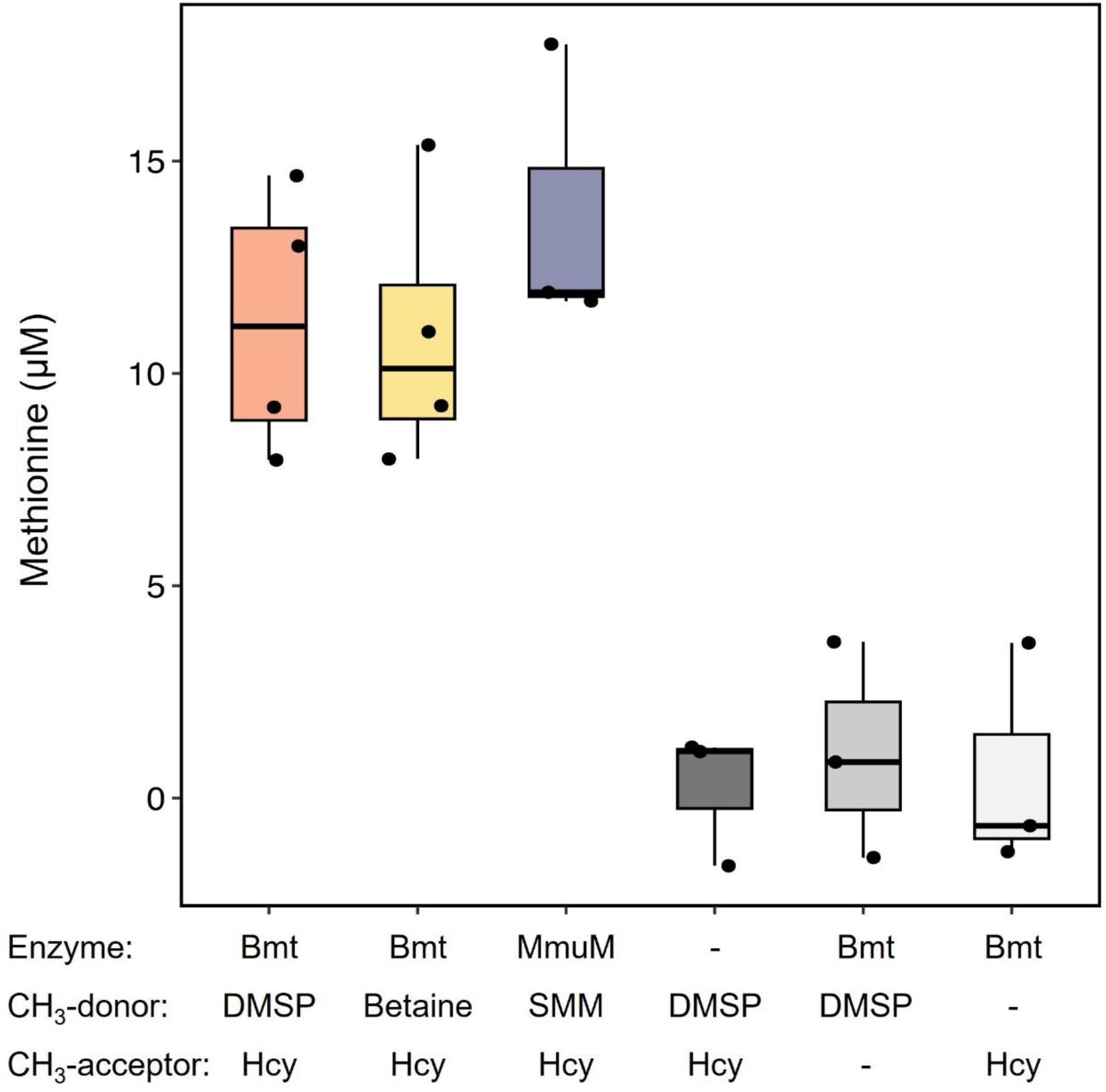
The *P. inhibens* Bmt utilizes DMSP and betaine as methyl group donors for coupled methionine synthesis. The formation of methionine was quantified using *in vitro* reactions containing purified Bmt enzyme together with the methyl group acceptor homocysteine (Hcy) and the methyl group donors DMSP or betaine. The purified methionine synthase MmuM of *E. coli* was used as a positive control, which is similar to Bmt and utilizes *S*-methylmethionine (SMM) as methyl group donor. Results were compared to negative control reactions in which single components were omitted (either Bmt, Hcy or DMSP). Box-plots show results from at least three independent experiments; black dots indicate individual measurements. Box-plot elements: center line - median; box limits - upper and lower quartiles; whiskers – min and max values.

### Inhibiting tetrahydrofolate synthesis reveals flexibility in methyl group assimilation required for lag phase shortening

To strengthen the observation that Bmt can act as a DMSP—homocysteine *S*-methyltransferase, potentially leading to methionine synthesis and lag phase shortening, our objective was to inhibit THF-dependent one-carbon metabolism in the cell. Such inhibition would disrupt THF-dependent DMSP demethylation by DmdA, as well as THF-dependent methionine synthesis by either the MetH-like split methionine synthase or the cobalamin-independent MetE enzyme (Fig. 1B). Importantly, inhibition of THF synthesis would not affect the DMSP—homocysteine *S*-methyltransferase activity of the stand-alone Bmt enzyme, which can function in a THF-independent manner (Fig. 3). Inhibition of THF synthesis was achieved by adding trimethoprim (TPM) to bacterial cultures^35^. Since TPM also inhibits THF-dependent nucleotide synthesis, cultures were supplemented with dNTPs to rescue bacterial growth (Fig. 4). Although bacterial growth was significantly disturbed by TPM, bacteria exhibited shorter lag phases in response to supplementation with 2 µm DMSP (Fig. 4). These findings suggest that bacteria possess an alternative route for methionine production, bypassing the THF-dependent steps. This alternative route could involve Bmt, as supported by the *in vitro* observation of DMSP—homocysteine *S*-methyltransferase activity (Fig. 3). To validate this hypothesis experimentally, it would be essential to inhibit THF synthesis in bacteria lacking Bmt. However, such validation presents challenges particularly with the *Δbmt* mutant, which is a methionine auxotroph requiring exogenous methionine supplementation. Therefore, while the potential of Bmt to couple DMSP demethylation with methionine synthesis was demonstrated *in vitro*, further investigation is needed to determine if Bmt performs similar functions within the bacterial cell.

**Figure 4.**
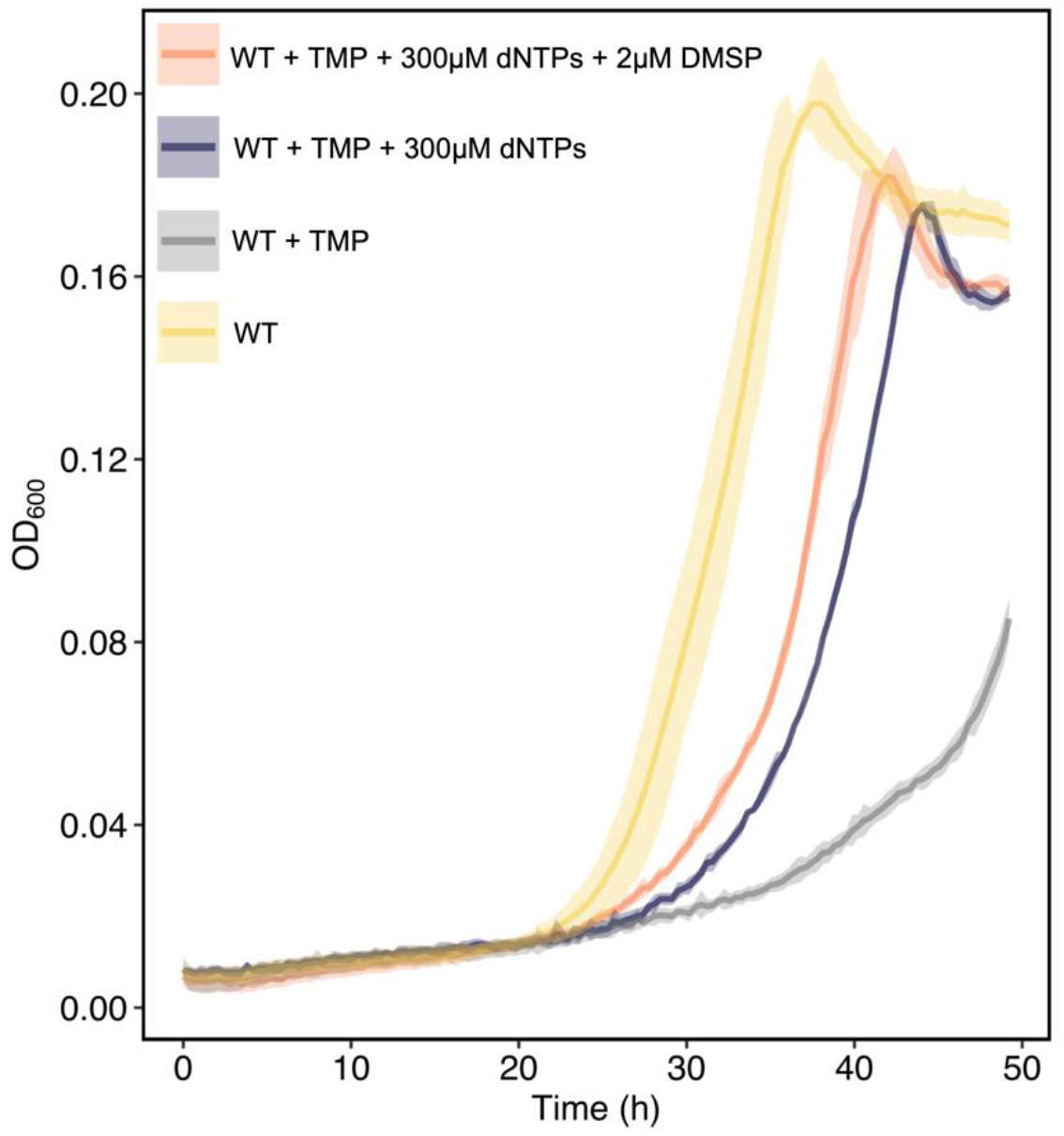
The co-factor tetrahydrofolate (THF) is not required for lag phase shortening with DMSP. THF synthesis was inhibited in *P. inhibens* cultures by adding 50 μg/ml trimethoprim (TMP). THF is required for nucleotide synthesis and therefore TMP-treated cells growth is affected (gray) in contrast with the control (yellow). Bacterial growth was rescued by the external addition of nucleotides (300 μM of dNTPs, purple and orange). Cultures treated with both TMP and dNTPs exhibited stimulated growth upon supplementation with 2 µM DMSP (orange) compared to control cultures without DMSP (purple). These results suggest that methyl groups can be harvested from DMSP and induce lag phase shortening in a THF-independent manner, possibly involving the THF-independent DMSP—homocysteine *S*-methyltransferase activity of Bmt. Lines represent the average growth curve based on three biological replicates and shaded areas indicate the standard deviation (SD).

### Lag phase shortening under stress conditions

Finally, we sought to gain insight into Bmt activity *in vivo*. Previously, we reported that a *P. inhibens ΔdmdA* mutant lacks the ability to harvest methyl groups from DMSP, which abolished DMSP-induced lag phase shortening, likely by preventing methyl group assimilation via the methionine cycle^3^. Following this observation, we now tested whether stress conditions could encourage the Bmt activity. Under stress conditions, DMSP methyl group assimilation by Bmt could become a co-factor independent and cost-effective alternative to the DmdA-MetH route. Therefore, cultures of the *ΔdmdA* mutant were exposed to high osmolarity (0.45 M NaCl) or oxidative stress (100 μM H_2_O_2_), these treatments stress bacteria as evident by markedly longer lag phases (Fig. 5). The cultures were supplemented with DMSP or betaine (2 μM), or untreated. Under non-stress conditions, the *ΔdmdA* mutant did not exhibit shorter lag times in response to DMSP, as previously reported^3^ (Fig. 5A). However, under high osmolarity and oxidative stress conditions, DMSP did trigger lag phase shortening in *ΔdmdA* mutant cultures (Fig. 5B-C). This observation suggests that Bmt could be an alternative route for DMSP methyl group assimilation by *P. inhibens*. Bmt may be particularly important under unfavourable conditions, a conclusion that requires further confirmation.

**Figure 5.**
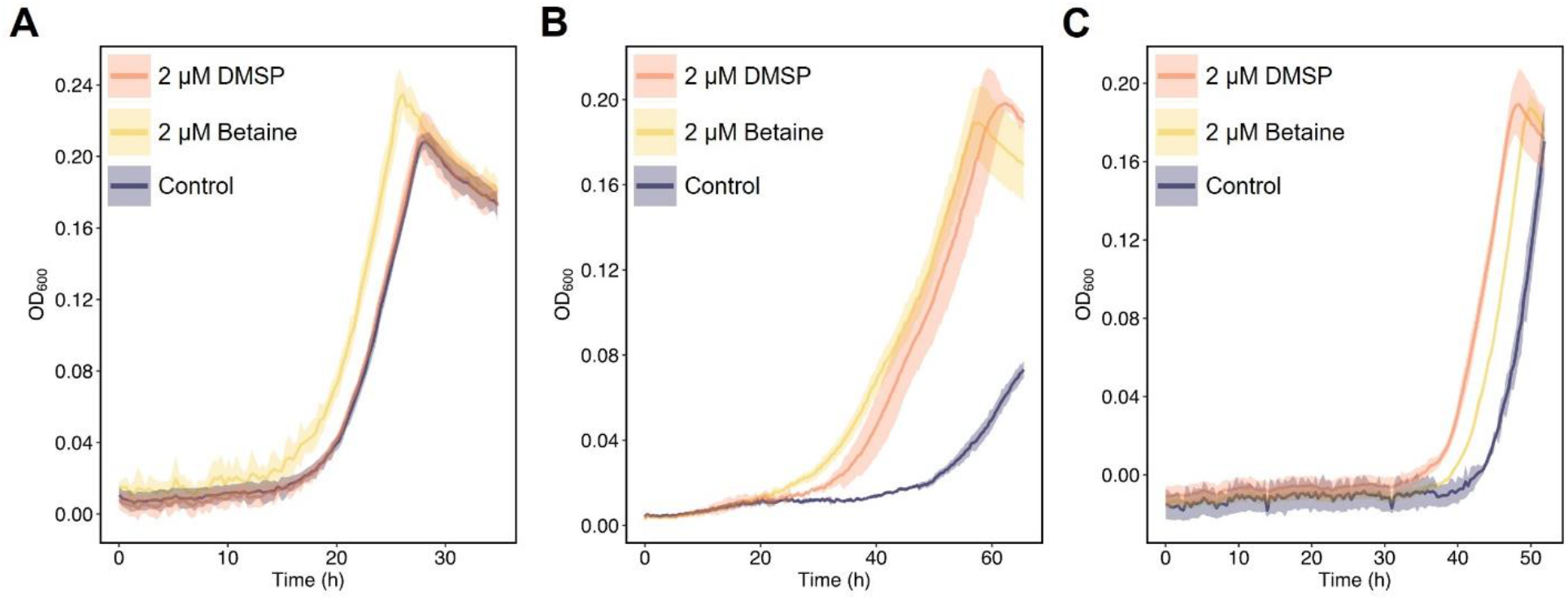
Lag phase shortening is triggered in the Δ*dmdA* mutant under high salt and oxidative stress conditions. **(A)** Under regular cultivation conditions in minimal media, the *ΔdmdA* mutant responds to betaine supplementation (2 µM, yellow), but not to DMSP (2 µM, orange), similar to the control (purple). However, under **(B)** high salinity conditions (0.45 M NaCl), and **(C)** oxidative stress (100 μM H_2_O_2_), DMSP does trigger lag phase shortening in the mutant (orange). Lines represent the average growth curve based on three biological replicates and shaded areas indicate the standard deviation (SD).

## Discussion

### The importance and versatility of methyl groups metabolism

Imbalances in methyl group metabolism, also termed C1 metabolism, are associated with human diseases^36^ and bacterial growth perturbations^6^. The major role of methyl group metabolism is to provide C1 groups for the synthesis of building blocks such as purines (adenine and guanine), thymine, methionine, histidine and polyamines^6,37^. Additional methyl groups are required to methylate DNA, RNA and proteins^38–40^. It was previously shown that the depletion of building blocks results in longer lag phases^2^, and that the synthesis of building blocks is upregulated during the lag phase of bacteria^41–44^. Deficiencies in macromolecule methylations can impair growth by influencing the cell cycle^45^, the circadian clock^46^ and chemotaxis proteins^47,48^. Importantly, the production of methyl groups is energetically costly. During this process, bacteria first produce a methylene group (involving the serine hydroxymethyltransferase and/or the glycine cleavage system^37,49^), which is then converted to a methyl group through the utilization of a NAD(P)H reducing equivalent^50^. This, and the requirement for additional co-factors, results in the fact that methionine—the only *S*-methylated amino acid—is the most expensive amino acid to produce^51^. It was shown that glucose-grown Roseobacter bacteria build the majority of their methionine from the methyl groups of supplemented DMSP^52^. Here, we show that methyl groups can be directly transferred from DMSP for the synthesis of methionine—a reaction that is catalyzed by Bmt (Fig. 3). These observations add versatility to the described routes of methionine synthesis^15,52^.

### Lag phase-related regulation of the methionine cycle

The methionine cycle is central in the regulation of the bacterial lag phase. Methionine is the only methylated compound that prolonged the lag phase of *P. inhibens*^3^. This effect can be either attributed to methionine directly, or to the accumulation of methionine cycle products such as homocysteine or S-adenosylmethionine (SAM). The methionine cycle is tightly feedback-regulated, explaining the absence of methionine overproducing bacteria outside the lab^53^. Similarly, imbalances in methionine synthesis are associated with diseases in higher organisms^54,55^. A negative feedback regulation was described for an *E. coli* enzyme that synthesizes homocysteine (MetA; homoserine *O*-succinyltransferase), showing that the enzyme is inhibited by methionine and SAM^25,56^. It was also demonstrated that methionine and/or homocysteine inhibit the growth of *E. coli* under certain conditions^24,57^. Moreover, the breakdown of methionine synthesis results in longer lag phases in *E. coli*^58^. In line with tight feedback-regulation of the methionine cycle, our data show that the methionine synthase gene *bmt* was downregulated in DMSP-supplemented bacteria during the lag phase (Fig. 1). Similarly, a homologues *bmt* gene was downregulated in a related Roseobacter bacterium when exposed to DMSP (accession: B5M07_08905^59^). In conclusion, we found that the methionine cycle plays a role in regulating the duration of the lag phase. Our data suggest that an elevated influx of methyl groups into this cycle can be counterbalanced by the downregulation of methionine synthase gene transcription. Results of our experiments with the *Δbmt* mutant support the centrality of the Bmt methionine synthase in lag phase shortening induced by methylated compounds.

### Bmt is advantageous under stress conditions

The possible importance of Bmt under conditions of osmotic pressure was previously discussed^13^. It was observed that a mutant strain of *S. meliloti* lacking the *metH* gene was unable to produce methionine, making it an auxotroph. However, when supplemented with 1 mM glycine betaine (GB), the mutant strain restored its growth, though at slower rates compared to methionine supplementation. This suggested an alternative pathway for methionine synthesis using GB as a methyl group donor. Further investigation revealed that this pathway is catalyzed by a betaine—homocysteine *N*-methyltransferase (Bmt), an enzyme previously characterized in humans and rats (BHMT)^60,61^. Interestingly, under high osmolarity conditions, such as exposure to 0.5 M NaCl, the addition of glycine betaine led to faster growth of the *metH* mutant compared to methionine supplementation alone^13^. While GB accumulation can be advantageous due to its osmoprotectant properties, these observations suggest that under hyper-osmotic conditions, Bmt can serve as an efficient demethylase of GB as well as a methionine synthase.

In addition, under high salinity conditions, addition of DMSP was reported to contribute to the fitness of *Vibrio* bacteria^62^. While there are no reported DMSP demethylases for *Vibrio* species, our previous observations demonstrated that *Vibrio* strains respond to methylated compounds^3^ and can shorten their lag phase. These data further support a possible link between methionine synthesis using DMSP as a methyl donor and salt stress conditions.

On the other hand, oxidative stress induces methionine auxotrophy in *E. coli* growing in minimal media, caused by the inactivation of the MetE enzyme and resulting in methionine limitation^63^. Additionally, *E. coli* cultures containing methionine resume growth faster after being exposed to oxidative stress conditions^63^. Our data demonstrate that under stress conditions, including high salt and oxidative stress, the *ΔdmdA* mutant is capable of expediting the lag phase using DMSP as a methyl group donor. The adaptability of the methionine cycle under stress conditions could provide an advantage for bacteria that constantly face changes in salinity and oxidative stress within the marine environment. Especially as these environmental challenges are intensified by anthropogenic activities^64^.

### Methionine synthesis in an environmental context

Methionine synthesis is primarily mediated by the cobalamin-dependent methionine synthase; however, its activity can be compromised by environmental factors like cobalamin deficiency or exposure to nitrous oxide^65,66^. Despite this knowledge, the specific environmental influences on bacterial methionine synthesis and the corresponding bacterial adaptation strategies remain largely unexplored.

Notably, while the metabolic pathway for methionine biosynthesis has been extensively characterized in *E. coli*, it is recognized that numerous bacterial species diverge from this canonical pathway. A comprehensive survey across bacterial species has underscored the non-canonical nature of the *E. coli* pathway, revealing a vast diversity in methionine synthesis mechanisms among bacteria^5^.

Additionally, microbial communities residing on algal cell surfaces, exemplified by the model marine bacterium *P. inhibens* in this study, contribute to the production of various compounds. This includes bacterial-produced methionine, which can significantly influence both the fitness of the algal host and the dynamics within the microbial community^67,68^. Thus, an intricate physiological interplay exists between microalgae and their associated bacteria, including the utilization of algal methylated compounds by bacteria to shorten their lag phase. In the complex web of microbial interactions, methionine synthesis emerges as a crucial metabolic process involved in shaping these relationships.

## Materials & Methods

### Key Resources Table

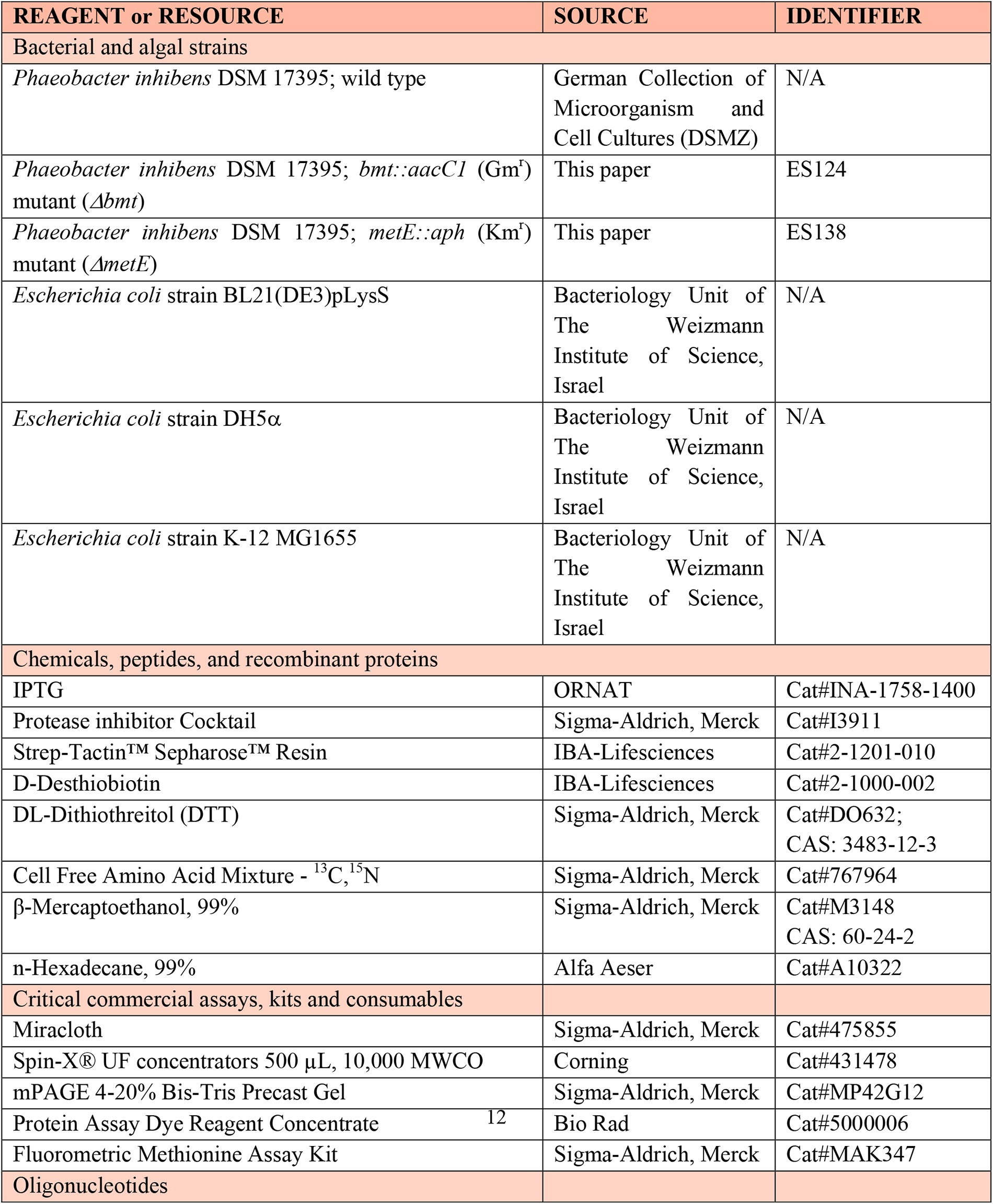
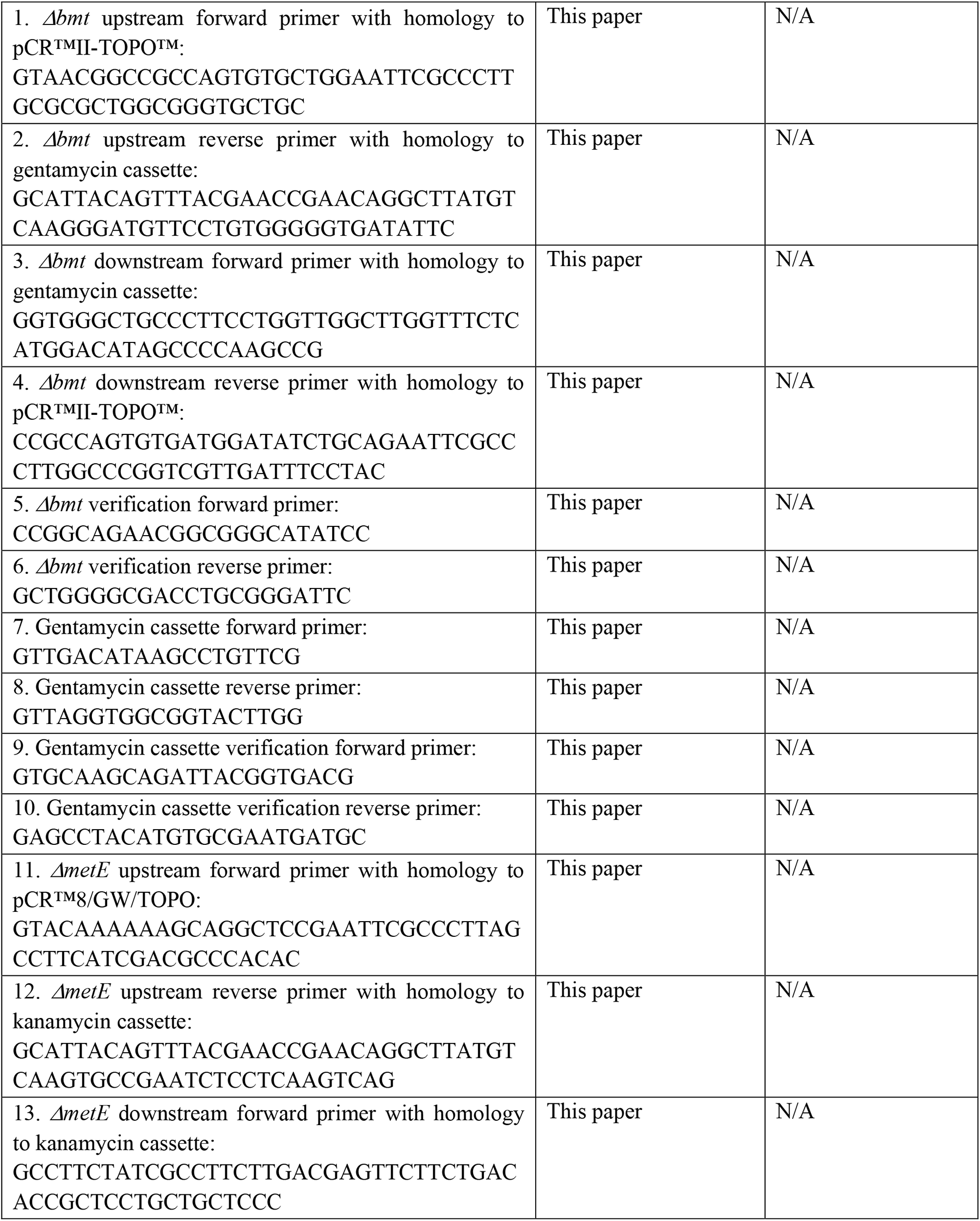
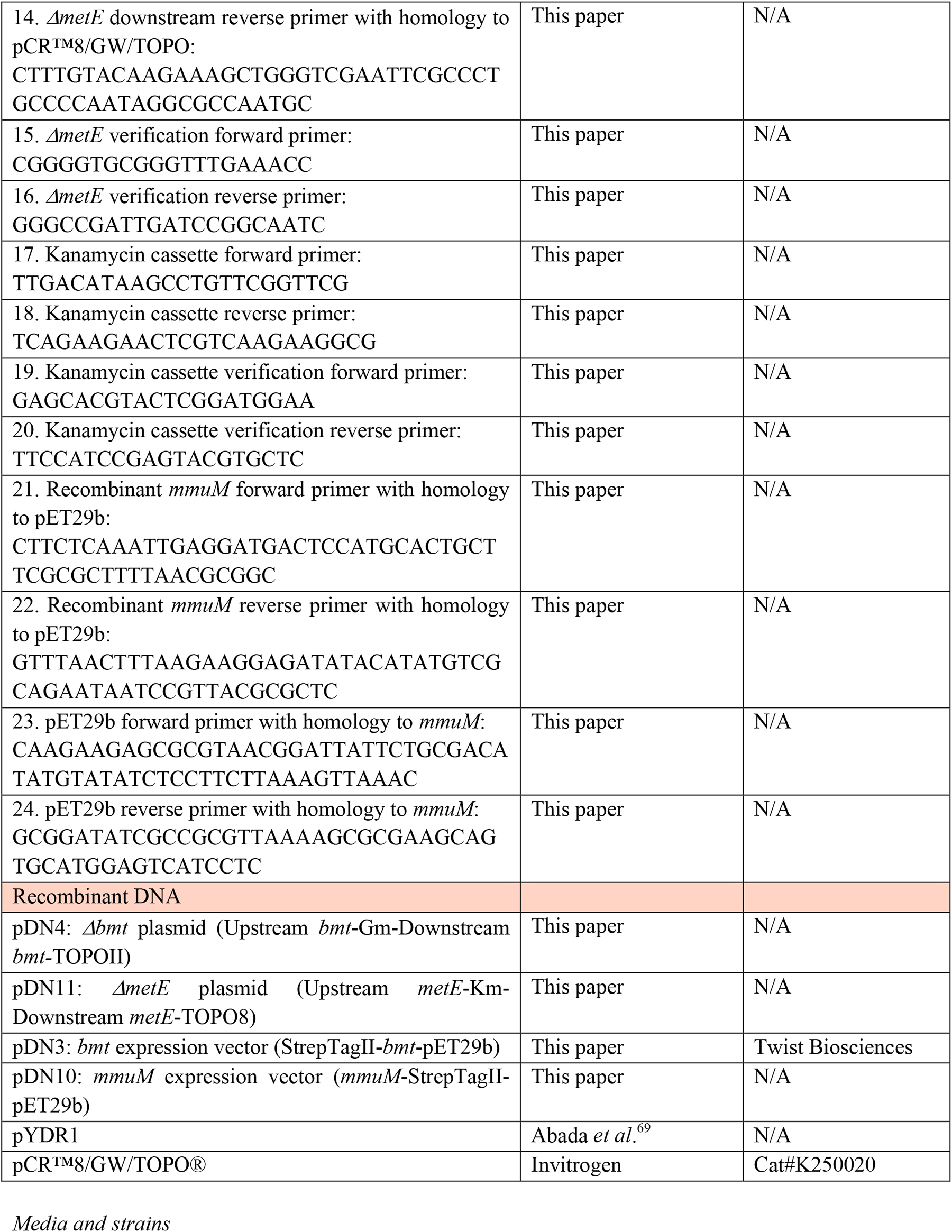

### Media and strains

The bacterial strain *Phaeobacter inhibens* DSM 17395 was purchased from the German Collection of Microorganism and Cell Cultures (DSMZ, Braunschweig, Germany). It was cultivated in ½ YTSS medium (yeast extract, 2 g/L; trypton, 1.25 g/L; sea salts; 20 g/L), or ASW medium based on the protocol of Goyet and Poisson^70^ and contained mineral salts (NaCl, 409.41 mM; Na_2_SO_4_, 28.22 mM; KCl, 9.08 mM; KBr, 0.82 mM; NaF, 0.07 mM; Na_2_CO_3_, 0.20 mM; NaHCO_3_, 2 mM; MgCl · 6 H_2_O, 50.66 mM; CaCl_2_, 10.2 mM, SrCl_2_ · 6 H_2_O, 0.09 mM), L1 trace elements (Na_2_EDTA · 2H_2_O, 4.36 mg/L; FeCl_3_ · 6 H_2_O, 3.15 mg/L; MnCl_2_ · 4 H_2_O, 178.1 μg/L; ZnSO_4_ · 7 H_2_O, 23 μg/L; CoCl_2_ · 6 H_2_O, 11.9 μg/L; CuSO_4_ · 5 H_2_O, 2.5 μg/L; Na_2_MoO_4_ · 2 H_2_O, 19.9 μg/L; H_2_SeO_3_, 1.29 μg/L; NiSO_4_ · 6 H_2_O, 2.63 μg/L; Na_3_VO_4_, 1.84 μg/L; K_2_CrO_4_, 1.94 μg/L), L1 nutrients (NaNO_3_, 882 μM; NaH_2_PO_4_ · 2 H_2_O, 36.22 μM), 5 mM NH_4_Cl, 33 mM Na_2_SO_4_, and 1 mM Glucose as carbon source, adjusted to a pH 8.2 with HCl. The same conditions were used for the mutants *Δbmt, ΔmetE and ΔdmdA,* with the addition of 30 μg/ml gentamycin, 150 μg/ml kanamycin and 30 μg/ml gentamycin respectively. Additionally, the mutant *Δbmt* was also supplemented with 200µM Methionine (Thermo Scientific). For high salinity experiments, 0.45 M NaCl (Bio-Lab Ltd, Jerusalem, IL) was added to the ASW medium, and for experiments with oxidative stress conditions 100 μM H_2_O_2_ (Invitrogen) was added to the ASW medium.

### Bacterial growth curves

Bacteria were streaked from glycerol stocks on ½ YTSS medium agar plates and incubated at 30 °C for 48 hours. After incubation, a single colony was used to inoculate a pre-culture with 10 mL ASW medium. The pre-culture was cultivated at 30 °C with 130 rpm shaking for 48 hours to reach stationary phase. Cells were then highly diluted (OD_600_ 0.00001) in fresh ASW, and supplemented, as indicated, with 2 μM DMSP, 2 μM betaine, 50 μg/ml Trimethoprim (TMP, Sigma-Aldrich, Merck, Darmstadt, Germany), 100 μM dNTPs (Thermo Scientific), 100 μM H_2_O_2_ (Invitrogen), or sterile water as control. Diluted cells were transferred to 96-well microtiter plate (150 µl per well) and overlaid with 50 µl hexadecane to prevent evaporation^71^. Bacterial growth was followed at 30°C in an Infinite 200 Pro M Plex plate reader (Tecan Group Ltd., Männedorf, Switzerland) with alternating cycles of 5 sec shaking and 19:55 min incubation. Absorption measurements were conducted at 600 nm following the shaking step and multiplied by a factor of 3.86 to reflect optical density measurements performed in 1 cm cuvettes.

### Lag phase RNA-sequencing

The RNA-sequencing dataset utilized in this study was previously reported by us^3^. Briefly*, P. inhibens* stationary cells were diluted to 0.01 in 30 mL of ASW media and supplemented with 50 µM DMSP or water as control and incubated at 30 °C with 130 rpm shaking. After 15 and 40 min of the inoculation, cells were collected by centrifugation at 4 °C. RNA extracts were obtained with the ISOLATE II RNA Mini Kit (Meridian Bioscience, OH, USA) and ribosomal rRNA depletion was performed using 100 pmol Pan-Bacteria probes. The obtained RNA-sequencing library was sequenced on a NextSeq 500 instrument with a 150 cycles Mid Output Kit (Illumina, San Diego, CA, USA) in paired-end mode. Quality filtered and trimmed sequencing reads were mapped to *P. inhibens* DSM 17395 genome (accession: GCF_000154765.2). DESeq2 was used for differential gene expression analysis by comparing DMSP-supplemented samples with control samples 15 min and 40 min after inoculation. Raw sequencing data were deposited under the BioProject accession PRJNA977030.

### P. inhibens Bmt and MetE null mutants

The primers and plasmids used for the KO constructs are described in the Key Resources Table. DNA manipulation and cloning PCR was performed using Phusion High Fidelity DNA polymerase (Thermo Scientific), according with manufacturer recommended PCR conditions. PCR-amplified DNA was cleaned with NucleoSpin Gel and PCR Clean-up kit (MACHEREY-NAGEL, Düren, Germany). Plasmid DNA was purified using QIAprep Spin Miniprep Kit (QIAGEN, Hilden, Germany).

For creation of *bmt* null mutant (*Δbmt*) cells (ES124), ∼1000 bp regions upstream and downstream of the *bmt* gene (accession: PGA1_c13370) were amplified by PCR, using the 1, 2, 3 and 4, respectively. The gentamycin resistance marker of pBBR1MCS5 was amplified using primers 7 and 8. The PCR-amplified fragments (upstream region + gentamycin resistance + downstream region) were assembled and cloned into pCR™II-TOPO™ vector (Invitrogen, Thermo Fisher Scientific, Waltham, MA, USA) using restriction-free cloning^72^, generating the plasmid pDN4.

For creation of *metE* null mutant (*ΔmetE*) cells (ES138), ∼1000 bp regions upstream and downstream of the *metE* gene (accession: PGA1_c27430) were amplified by PCR, using the primers 11, 12, 13 and 14 respectively. The kanamycin resistance marker of pYDR1 was amplified using primers 17 and 18. The PCR-amplified fragments (upstream region + kanamycin resistance + downstream region) were assembled and cloned into the pCR™8/GW/TOPO® vector (Invitrogen) using restriction-free cloning^72^, generating the plasmid pDN11.

*P. inhibens* electrocompetent cells (300 μl) were transformed with 10 μg of the constructed plasmids by a pulse of 2.5 kV (Bio Rad), and cells were selected on ½ YTSS plates containing 30 μg/ml Gentamycin or 150 μg/ml Kanamycin. Successful null mutants were verified in single cell clones by PCR (5, 6, 9, 10 for *Δbmt* KO, and 15, 16, 19, 20 for *ΔmetE*) and sanger sequencing.

### Methionine synthesis with purified Bmt

Primers and plasmids used for heterologous expression of Bmt and MmuM (positive control) are listed in the Key Resources Table. To produce *P. inhibens* Bmt (accession: PGA1_c13370), a pET29b expression vector was designed that encodes the *bmt* gene together with an N-terminal Strep-tag II peptide sequence (pDN3). The vector was synthesized by Twist Biosciences (San Francisco, CA, USA). To produce *E. coli* MmuM (accession: Q47690), the respective gene was PCR-amplified from *E. coli* K-12 MG1655 (primers 27-28). The expression vector was PCR-amplified from pET29b (primers 29-30) and cloned together with the *mmuM* PCR product using the CPEC technique^73^. This resulted in a MmuM expression vector with C-terminally fused Strep-tag II (pDN10). Plasmid construction was validated by Sanger sequencing. *E. coli* BL21 competent cells were transformed by electroporation with 100 µg expression vector, using a pulse of 1.8 kV (MicroPulser, Bio-Rad Laboratories, Hercules, CA, USA). Electroporated cells were selected for positive transformants using LB medium agar plates with 50 µg/ml kanamycin.

For protein expression, *E. coli* BL21 transformants were grown in TYG medium with 50 µg/ml kanamycin at 37°C until reaching the mid-log phase (OD_600_ = 0.7). Protein expression was induced by adding 0.2 mM IPTG. Cells were harvested three hours after IPTG induction by centrifugation (4,000 rpm, 15 min, 4 °C). Cell pellets were resuspended in 20 mL NP buffer (50 mM NaH_2_PO_4_, 300 mM NaCl, pH 8) with 100 µl 10X Protease Inhibitor Cocktail (Sigma-Aldrich, Merck, Darmstadt, Germany). Resuspended cells were filtered with Miracloth (Sigma-Aldrich, Merck, Darmstadt, Germany) and passed three times through a French Pressure cell press (15,000 psi) for cell disruption. Disrupted cells were centrifuged (4,000 rpm, 15min, 4 °C) and the supernatant was loaded onto a Strep-Tactin Sepharose resin column (IBA-Lifesciences, Göttingen, Germany). The column was washed three times with 5 mL NP buffer, and proteins were eluted with 3 mL NPD buffer (50 mM NaH_2_PO_4_, 300 mM NaCl, 2.5 mM Desthibiotin, pH 8). Eluted proteins were concentrated with a 10 kDa membrane (Spin-X® UF 500 µL Centrifugal Concentrator, 10,000 MWCO; Corning, NY, USA) in 20 mM HEPES-KOH buffer (pH 7.5). Protein concentrations were determined using the Protein Assay Dye Reagent Concentrate (Bio-Rad Laboratories, Hercules, CA, USA). The purity was controlled by electrophoresing 3 µg of protein with SDS-sample buffer on a polyacrylamide gel (mPAGE™ 4-20% Bis-Tris Precast Gel, Merck, Darmstadt, Germany), followed by Bio-Safe Coomassie staining (Bio-Rad Laboratories, Hercules, CA, USA; fig. S14).

Methionine synthesis with either DMSP or betaine as methyl group donors was measured *in vitro*, using purified enzyme and a protocol adapted from Ranocha *et al*.^74^. The *in vitro* reactions contained buffer (20 mM HEPES-KOH, pH 7.5, 2 mM dithiothreitol), a methyl group acceptor (2 mM homocysteine), a methyl group donor (200 µM of either DMSP, betaine or SMM) and purified enzyme (200 µM of either Bmt or MmuM). The *in vitro* reactions (v = 50 µl) were incubated for 2 hours at 30 °C. Methionine synthesis was measured using the Methionine Assay Kit (Sigma-Aldrich, Merck, Darmstadt, Germany). For measurements, 20 µl of each *in vitro* reaction was mixed with 30 µl Met Assay Buffer and 50 µl Met Assay Reaction Mix. In parallel, 20 µl of the same *in vitro* reactions were mixed with 30 µl Met Assay Buffer and 50 µl Met Assay Background Control Mix. The mixes were incubated for 30 min at 37 °C, followed by fluorescence readings (λ_ex_ = 535 nm/λ_em_ = 587 nm; Infinite 200 Pro M Plex plate reader, Tecan Group Ltd., Männedorf, Switzerland). The Methionine Assay Kit includes a methionine standard for absolute quantifications. The steps needed to produce purified proteins and to measure enzymatic methionine formation were repeated in three independent experiments.

## Data availability

This study includes no data deposited in external repositories. The RNA-sequencing source data utilized in this study was previously deposited under the BioProject accession ID PRJNA77030.^3^

## Acknowledgments

D.A.N.B received the Armando and Maria Jinich Fellowship. M.S. received a Dean of Faculty Fellowship, a Sir Charles Clore Fellowship (Clore Israel Foundation) and a Senior Postdoc Fellowship. The study was funded by the European Research Council (ERC StG 101075514) and the de Botton center for marine sciences, granted to E.S.

## Author contributions

D.A.N.B., M.S., and E.S. designed the study and wrote the manuscript. D.A.N.B. and M.S. performed and analyzed experiments.

